# Palmitoylation regulates norepinephrine transporter trafficking and expression and is potentially involved in the pathogenesis of postural orthostatic tachycardia syndrome

**DOI:** 10.1101/2024.03.22.586171

**Authors:** Christopher R. Brown, James D. Foster

## Abstract

Postural orthostatic tachycardia syndrome (POTS) is an adrenergic signaling disorder characterized by excessive plasma norepinephrine, postural tachycardia, and syncope. The norepinephrine transporter (NET) modulates adrenergic homeostasis *via* reuptake of extracellular catecholamines and is implicated in the pathogenesis of adrenergic and neurological disorders. Previous research has outlined that NET activity and trafficking is modulated *via* reversible post-translational modifications like phosphorylation and ubiquitylation. S-palmitoylation, or the addition of a 16-carbon saturated fatty acid, is another post-translational modification responsible for numerous biological mechanisms. In this study, we reveal that NET is dynamically palmitoylated and inhibition of this modification with the palmitoyl acyltransferase (DHHC) inhibitor, 2-bromopalmitate (2BP), results in decreased NET palmitoylation within 90 min of treatment. This result was followed closely with a reduction in transport capacity, cell surface, and total cellular NET expression after 120 min of treatment. Increasing 2BP concentrations and treatment time revealed a nearly complete loss of total NET protein. Co-expression with individual DHHCs revealed a single DHHC enzyme, DHHC1, promoted WT hNET palmitoylation and elevated NET protein levels. The POTS associated NET mutant, A457P, exhibits dramatically decreased transport capacity and cell surface levels which we have confirmed in the current study. In an attempt to recover A457P NET expression we co-expressed the A457P variant with DHHC1 to drive expression as seen with the WT protein but instead saw an increase in NET N-terminal immuno-detectable fragments. Further investigation of A457P NET palmitoylation and surface expression is necessary, but our preliminary novel findings reveal palmitoylation as a mechanism of NET regulation and suggest that dysregulation of this process may contribute to the pathogenesis of POTS.

## INTRODUCTION

Norepinephrine (noradrenaline [NE]) is a monoamine neurotransmitter involved in modulating the body’s neuro-cognitive and sympathetic processes. The sympathetic nervous system (SNS) is a division of the autonomic nervous system that controls the body’s “fight or flight” response. Sympathetic outflow originates from nuclei in the *rostral ventrolateral medulla* (RVLM), where it projects inferiorly within the spinal cord and extends to pre-ganglionic neurons in the thoracolumbar region (1,2). Subsequently, stimulation of postganglionic neurons releases NE, where it diffuses and modulates various effector-tissue adrenergic receptors to regulate digestion, metabolism, and cardiovascular functions like heart rate, contractility, and vascular tone (3). Noradrenergic homeostasis is tightly regulated *via* reuptake of NE by the Na^+^/Cl^-^ dependent norepinephrine transporter (NET). Following re-uptake, NE is re-packaged into pre-synaptic vesicles where it is prepared for release. The abundance of physiologic functions, and pathophysiologic implications, mediated by NE signaling makes NET a target of interest in biomedical research and therapeutic design (4–7). NET activity and surface expression has been found to be abnormal in cardiovascular diseases including hypertension, cardiomyopathy, heart failure, and myocardial ischemia, (8–12). Therapeutically, NET is a target for prescription and illicit chemical substances like tricyclic anti-depressants (TCA), serotonin-norepinephrine reuptake inhibitors (SNRIs), cocaine, and amphetamines, which all act to transiently elevate the extracellular concentration of NE either through direct blockade or substrate-mediated efflux.

In normal physiologic conditions, about 750 ml of thoracic blood is transferred downward when an individual stands from a seated position. This sudden drop in blood pressure is sensed by baroreceptors of the carotid sinus and aortic arch, which potentiate signals to the *nucleus tractus solitarius* (NTS) by way of the glossopharyngeal (CNIX) and vagus (CNX) nerves (1,2,13,14). The NTS relays this signal to the RVLM, and then the cardiovascular system *via* efferent sympathetic nerves to remedy the gravity-induced hypotension. In other words, transient orthostatic stress activates the SNS to maintain cerebral perfusion by increasing heart rate, cardiomyocyte contractility, and vascular smooth muscle contraction.

Commonly occurring in women between the ages of 20-50, postural orthostatic tachycardia syndrome (POTS), also known as orthostatic intolerance (OI), is a syndrome of excessive sympathetic activity triggered by orthostatic stress (15,16). With no evidence for changes in sympathetic outflow, this state of hyperadrenergia is characterized by symptoms of postural tachycardia, syncope, and elevated levels of plasma NE without significant orthostatic hypotension (15–19). Investigations into the pathogenesis of POTS have suggested there is a dysfunction in NET, identifying characteristics like decreased NE clearance, increased spill-over, and the absence of response to tyramine (19–21). The oxidized intraneuronal metabolite of NE, dihydroxyphenolglycol, is reported as low in relation to the concentration of NE, suggesting a defect in reuptake by NET and subsequent intra-neuronal metabolism (16,18). Likewise, in human studies, NET blockade from cocaine, amphetamines, and TCAs produce the characteristic symptoms of POTS (16,22–24).

Efforts to uncover a molecular basis for POTS have identified a rare mutation, A457P, in the 9^th^ transmembrane domain of NET (16). Twin patients who were heterozygous for A457P NET demonstrated classic symptoms of POTS, including a chronic history of exertional and orthostatic tachycardia, dyspnea, and syncope (16). *In vitro* analysis of A457P NET revealed A457P to be a dominant negative mutation, with impaired biogenesis driving a >98% loss in transport capacity (17). Hahn and colleagues have successfully generated A457P NET knock-in mice and analysis of this model indicated that NET has a lowered transport capacity, surface expression, and total cellular NET density that generated the characteristic phenotype of POTS (18). Consistent with these findings, post-mortem analysis of patients with POTS revealed a large decrease in total postganglionic NET expression, supporting the hypothesis that dysfunction of NET biogenesis and/or degradation is a component of the pathogenic mechanism (25).

Similar to the process of phosphorylation, S-palmitoylation is dynamic and reversible, permitting physiologic control over protein folding, trafficking, localization, and activity in accordance with cellular demands (26–28). Palmitoylation has been identified to regulate dopamine transporter (DAT) kinetics and stability (26,29), serotonin transporter (SERT) kinetics, trafficking, and expression (30), and biogenesis of the cystic fibrosis transmembrane regulator (CFTR) (31). The diverse range of targets and functional consequences controlled by palmitoylation emphasizes its importance in maintaining normal cellular and physiologic homeostasis. In the current study, we found that NET is regulated by S-palmitoylation, and that inhibition of this process alters NET surface levels and total cellular expression. We also found that inhibiting NET palmitoylation has similar outcomes when compared with the A457P coding variant, suggesting that palmitoylation may be necessary in the maturation and expression of NET, contributing to the biogenesis of NET and potentially the pathogenesis of POTS.

## MATERIALS AND METHODS

### Materials

[phenyl 2,5,6-^3^H]DA (>30 Ci/mmol) was from Vitrax; DA and hydroxylamine were from Sigma Millipore; DAT monoclonal antibody 16 (MAb 16) was previously authenticated (32), anti-mouse NET-05 and anti-human NET17-1 antibodies were from Mab Technologies. N-ethylmaleimide (NEM), high capacity NeutrAvidin-agarose resin, and bicinchoninic acid protein assay reagent were from Thermo Scientific. Sulfo-NHS-SS-biotin was from ProteoChem. HPDP-biotin was from Apex Bio. Rats were purchased from Envigo. All animals were housed and treated in accordance with regulations established by the National Institutes of Health and approved by the University of North Dakota Institutional Animal Care and Use Committee.

### Cell Culture

Lilly Laboratory Cell Porcine Kidney (LLC-PK_1_) cells stably expressing WT hNET or A457P hNET were grown in Dulbecco’s Modified Eagle Medium (DMEM) containing 5% fetal bovine serum, 100 μg/mL penicillin/streptomycin, and supplemented with 400 μg/mL of Geneticin (G418) for maintenance of stable expression. Cells were maintained in a humidified incubator gassed with 5% CO_2_ at 37^°^C. NET expression levels were verified by sodium dodecyl sulfate-polyacrylamide gel electrophoresis (SDS-PAGE) and immunoblotting of the cellular lysates against anti-human NET specific antibody (NET17-1 – MAb Technologies).

### Transfection

LLC-PK_1_ cells stably expressing WT hNET or A457P hNET were seeded into 24-well (0.5 µg DNA/well) or 100 mM plates (5 µg DNA/plate) and grown to 80% confluence. Cells were then transfected with the indicated amount of the appropriate plasmid using X-tremeGENE HP reagent (Sigma Millipore) and experiments were performed 24 hours post-transfection.

### Membrane Preparation

LLC-PK_1_ cells expressing the indicated NETs were grown in 100 mm plates to 85% confluency. Cells were washed twice with 3 mL of ice-cold Buffer B (0.25 mM sucrose, 10 mM triethanolamine, pH 7.8), scraped, and collected in 500 µL of Buffer B containing a protease inhibitor cocktail of 1 µM phenylmethylsulphonyl fluoride (PMSF) and 5 µM ethylenediaminetetraacetic acid (EDTA) at 4^°^C and transferred to a 2 mL microcentrifuge tube. Cells were then pelleted *via* centrifugation at 3,000 *x g* for 5 min at 4^°^C, the supernatant fraction was removed, and the cell pellet was suspended in 1 mL of ice-cold Buffer C (0.25 M sucrose, 10 mM triethanolamine, 1 mM EDTA, 1 µM PMSF, pH 7.8) and subsequently homogenized *via* 30 strokes of the pestle in a Dounce homogenizer. Homogenates were cleared of cellular debris and nuclei by centrifugation at 800 *x g* for 10 min. The post-nuclear supernatant fraction was collected and centrifuged at 18,000 *x g* for 12 min at 4°C to pellet cell membranes. The resulting membrane pellet was suspended in 1 mL of sucrose phosphate (SP) buffer (10 mM sodium phosphate, 0.32 M sucrose, pH 7.4 with 1 μM PMSF and 5 μM EDTA) and assayed for protein concentration.

### Synaptosome Preparation

Male Sprague-Dawley rats weighing between 200 and 300 g were decapitated, and striatal hemispheres were excised, weighed, and placed in ice-cold SP buffer. The striata were homogenized in 2 mL ice-cold SP buffer with 30 up and down strokes in a glass-Teflon homogenizer, diluted further with 8 mL SP buffer, mixed and centrifuged at 3000 *x g* for 2 min at 4^°^C. The supernatant fraction was isolated and subjected to an additional centrifugation at 17,000 *x g* for 12 min. The resulting P2 pellet encompassing synaptosomes was suspended in SP buffer containing protease inhibitors to 20 mg/mL using the original wet weight of the striatal tissue.

### Acyl-Biotinyl Exchange (ABE)

The ABE method used as previously described (30) was adapted from Wan et. al (33) where palmitoylated proteins are detected in three steps: (I) Free cysteine thiols are blocked; (II) thioester linked palmitoyl groups are removed by hydroxylamine (NH_2_OH); (III) the formerly palmitoylated and now newly generated sulfhydryl groups are biotinylated. Membranes prepared from LLC-PK_1_ cells expressing WT hNET were solubilized in 250 μL of lysis buffer (50 mM HEPES pH 7.0, 2% SDS (w/v), 1 mM EDTA) containing 25 mM N-ethylmaleimide (NEM), incubated for 20 min in a 37^°^C water bath and mixed end-over-end for at least 1 hour at ambient temperature. Proteins were precipitated *via* the addition of 1 mL acetone and centrifugation at 18,000 *x g* for 10 min. The protein pellet was resuspended in 250 µl lysis buffer containing 25 mM NEM and incubated for 1 hour at ambient temperature. This process was repeated a final time with incubation overnight with end-over-end mixing. NEM was removed *via* acetone precipitation/centrifugation and the protein pellet was resuspended in 250 μL 4SB buffer (50 mM Tris, 5 mM EDTA, 4% SDS, pH 7.4) and residual NEM was removed by an additional acetone precipitation and centrifugation. The pellet was resuspended in 200 μL 4SB buffer and thioesterified palmitate molecules were removed by incubation with hydroxylamine (NH_2_OH) where the sample was split into two equal aliquots (100 μL) with one diluted with 800 µl Tris-HCl, pH 8.0 (negative control) and the other diluted with 800 µL NH_2_OH (0.7 M final) and incubated at ambient temperature for 30 min with end-over-end mixing. Both samples were treated with 100 μL of a sulfhydryl-specific biotinylating reagent, HPDP-biotin (0.4 M final concentration) and incubated at ambient temperature with end-over-end mixing for 1 h. NH_2_OH and biotin reagents were removed by acetone protein precipitation with centrifugation at 18,000 *x g* and supernatant fraction aspiration. Protein pellets were then resuspended and solubilized in 150 μL 4SB, proteins precipitated with 600 µL of acetone, with centrifugation at 18,000 *x g* followed by supernatant fraction aspiration. The final pellet was suspended in 75 μL ABE lysis buffer and a 10 μL aliquot was set aside for total NET determination by immunoblotting while 65 μL was diluted with 1500 μL Tris buffer and incubated with 50 μL of a 50% slurry of Neutravidin resin overnight at 4^0^C with end-over-end rotation to affinity purify the biotinylated proteins/peptides. Unbound proteins/peptides were washed away by three cycles of 8,000 *x g* centrifugation, supernatant fraction aspiration and resuspension with 750 μL radio immunoprecipitation assay buffer (RIPA: 1% Tx-100, 1% sodium deoxycholate, 0.1% SDS, 125 mM sodium phosphate, 150 mM NaCl, 2 mM EDTA, 50 mM NaF). Proteins and/or peptides were eluted from the resin by incubation in 2x Laemmli SB for 20 min at ambient temperature. Samples were then subjected to SDS-PAGE, transferred to polyvinylidene fluoride (PVDF) and immunoblotted with mouse anti-human NET (NET17-1).

### Cell Surface Biotinylation

LLC-PK_1_ cells stably expressing WT hNET were grown in 24-well plates until 80% confluent. The cells were washed twice with 0.5 mL 37^°^C Krebs Ringers HEPES (KRH: 120 mM NaCl, 5 mM KCl, 2 mM CaCl, 1 mM MgCl, 25 mM NaHCO_3_, 5.5 mM HEPES, 1 mM D-Glucose) buffer and re-incubated in KRH buffer with or without 7.5 μM 2BP for 30, 60, 90, and 120 min at 37^°^C. Immediately following treatment, the cells were washed three times with ice-cold Hank’s balanced salt solution containing Mg^2+^ and Ca^2+^ ions (HBSS Mg-Ca: 1 mM MgSO_4_, 0.1 mM CaCl_2_, pH 7.4), and subsequently incubated twice with 0.5 mg/mL of membrane-impermeable sulfo-NHS-SS-biotin for 25 min on ice with rocking. The biotinylation reagent was removed and the reaction was quenched by two sequential incubations with 100 mM glycine in HBSS Mg-Ca for 20 min on ice with rocking. Cells were washed with HBSS Mg-Ca and lysed with 125 μL/well RIPA buffer containing protease inhibitor and the lysates from 12 wells were pooled in a 2 mL microcentrifuge tube. Protein (50 μg) from each lysate was incubated with 50 µL of a 50% slurry of NeutrAvidin resin overnight at 4°C with end-over-end rotation. The beads were washed three times with RIPA buffer, and the bound protein was eluted with 32 μL of SB followed by SDS-PAGE and immunoblotting for NET with anti-human NET antibody (NET 17-1).

### [^3^H]DA Uptake Assay

LLC-PK_1_ cells stably expressing WT hNET were grown in 24-well plates until 80% confluent and were treated with indicated concentrations of 2BP prepared in DMSO for 30-minute intervals at 37 °C followed by [^3^H]DA uptake assay in KRH. Final DMSO concentration was 1% or less, which by itself did not affect DA transport activity, and equal DMSO concentrations were used in all vehicle controls. Immediately following 2BP incubation, NET uptake was conducted for 8 min at 37^°^C in triplicate with 10 µM total DA containing 10 nM [^3^H]DA; nonspecific uptake was determined by addition of 100 µM (-)cocaine. Following uptake, the cells were rapidly washed three times with 500 µL of ice-cold KRH buffer and then solubilized in 1% Triton X-100 for at least 20 min, and radioactivity contained in lysates was assessed by liquid scintillation counting with uptake values being normalized to total cellular protein (pmol/min/mg).

### Saturation Analysis

LLC-PK_1_ cells stably expressing WT hNET were grown in 24-well plates until 80% confluent. The cells were washed with 0.5 mL 37^°^C KRH buffer and re-incubated in KRH buffer with or without 7.5 μM 2BP for 30 min at 37^°^C. Immediately following 2BP incubation, NET uptake was conducted in triplicate for 8 min at 37^°^C with 0.32, 0.6, 1, 3, 10, and 20 μM total DA containing 10 nM [^3^H]DA; nonspecific uptake was determined in the presence of 100 µM cocaine. Cells were rapidly washed twice with 500 µL ice-cold KRH buffer and lysed with 1% Tx-100 for at least 20 min with rocking at ambient temperature. Lysates were collected and analyzed by liquid scintillation counting. Kinetic values were determined using Prism software and nonlinear regression analysis, and V_max_ values were normalized to total cellular protein (pmol/min/mg) and transporter surface levels determined by surface biotinylation assays performed in parallel for each experiment.

### SDS-PAGE and Immunoblotting

Proteins were denatured in SB and were electrophoretically resolved using 4-20% polyacrylamide gels. Proteins were then transferred onto PVDF membrane and immunoblotted for NET using anti-human NET (NET17-1) antibody diluted 1:1,000 in blocking buffer (3% bovine serum albumin, phosphate buffered saline (PBS: 137 mM NaCl, 2.7 mM KCl, 10 mM NaH_2_PO_4_, 1.8 mM KH_2_PO_4_) and incubated for at least two hours at ambient temperature or overnight at 4°C. After five washes for 5 min with PBS containing 0.1% Tween, the membrane was incubated for 45 min in alkaline phosphatase (AP)-linked anti-mouse IgG secondary antibody diluted 1:5,000 in blocking buffer followed by 5 additional 5 min washes with 0.1% Tween/PBS. Protein bands were visualized by chemiluminescence using Immun-Star™ AP substrate (Bio-Rad) applied to the membrane and incubated for 5 min at ambient temperature. Band intensities were quantified using Quantity One® software (Bio-Rad).

## RESULTS

### Identification of NET Palmitoylation and Inhibition by 2BP

In previous studies we revealed that DAT is palmitoylated, and have characterized its role in controlling DAT kinetics, stability, and its reciprocal relationship with DAT phosphorylation (26,29,34). In this study, we extended our examination of palmitoylation in monoamine transporters to include NET. The similarity in transporter sequence homology and catecholamine physiology, stirred our interest to investigate the potential similarities and differences in how palmitoylation may impact NET regulation. In our initial experiments, we used LLC-PK_1_ cells stably expressing WT hNET and rat brain synaptosomes prepared from locus coeruleus, hippocampus, thalamus, and striatum. Our selection of LLC-PK_1_ cells as the primary heterologous system was based on our previous successes in characterizing DAT (26,29,34) and SERT (30) palmitoylation in this heterologous system. Membranes or synaptosomes prepared from these cells or rat brain regions, respectively, underwent acyl-biotinyl exchange (ABE) analysis followed by SDS-PAGE and immunoblotting with either anti-human NET (NET-17-1), anti-mouse NET (NET-05), or anti-rat DAT (Mab16) murine monoclonal antibodies.

These data revealed that NET is palmitoylated in both native rat tissue and LLC-PK_1_ cells. In the hydroxylamine (NH_2_OH) treated fraction of Figure 1A, a modest band running at ∼90 kDa was observed in rat synaptosomes when blotted with NET-05 Ab, suggesting a thioesterified version of rNET. This finding was consistent with blotting the same samples for DAT (Mab-16), confirming palmitoylation of rDAT in the striatum fraction (*Fig. 1B*). Importantly, the same bands were not evident in the Tris fraction, verifying the lack of biotin incorporation in this negative control for the ABE assay resulting in minimal nonspecific NET pull-down and detection. Interestingly, the 90 kDa band was minimal in the total fractions of both NH_2_OH and Tris treated samples (*Fig. 1A*). In these fractions, a stronger band at ∼60 kDa was present that likely represents the immature unglycosylated or core glycosylated form of rat NET (rNET) (*Fig. 1A & B, IM*). Following the successful identification of palmitoylated rNET in native tissue, we subjected WT hNET expressed in LLC-PK_1_ cells to ABE analysis. A strong band at ∼90 kDa was observed in the ABE and total samples when blotted with NET-17-1 Ab (*Fig. 1C*) and was absent in the control (Tris) of the ABE (*Fig. 1C, ABE upper panel*) and in parental LLC-PK_1_ cells which do not express NET (*Fig. 1C*, ABE and IB in far-left lane). The lower immunoblot panel (IB) represents the total NET level in aliquots taken just prior to neutravidin pull-down and is used to normalize the quantification of palmitoylation in the upper panels (*ABE*) where palmitoylated NET is a fraction of the total NET present in the sample.

**FIGURE 1.**
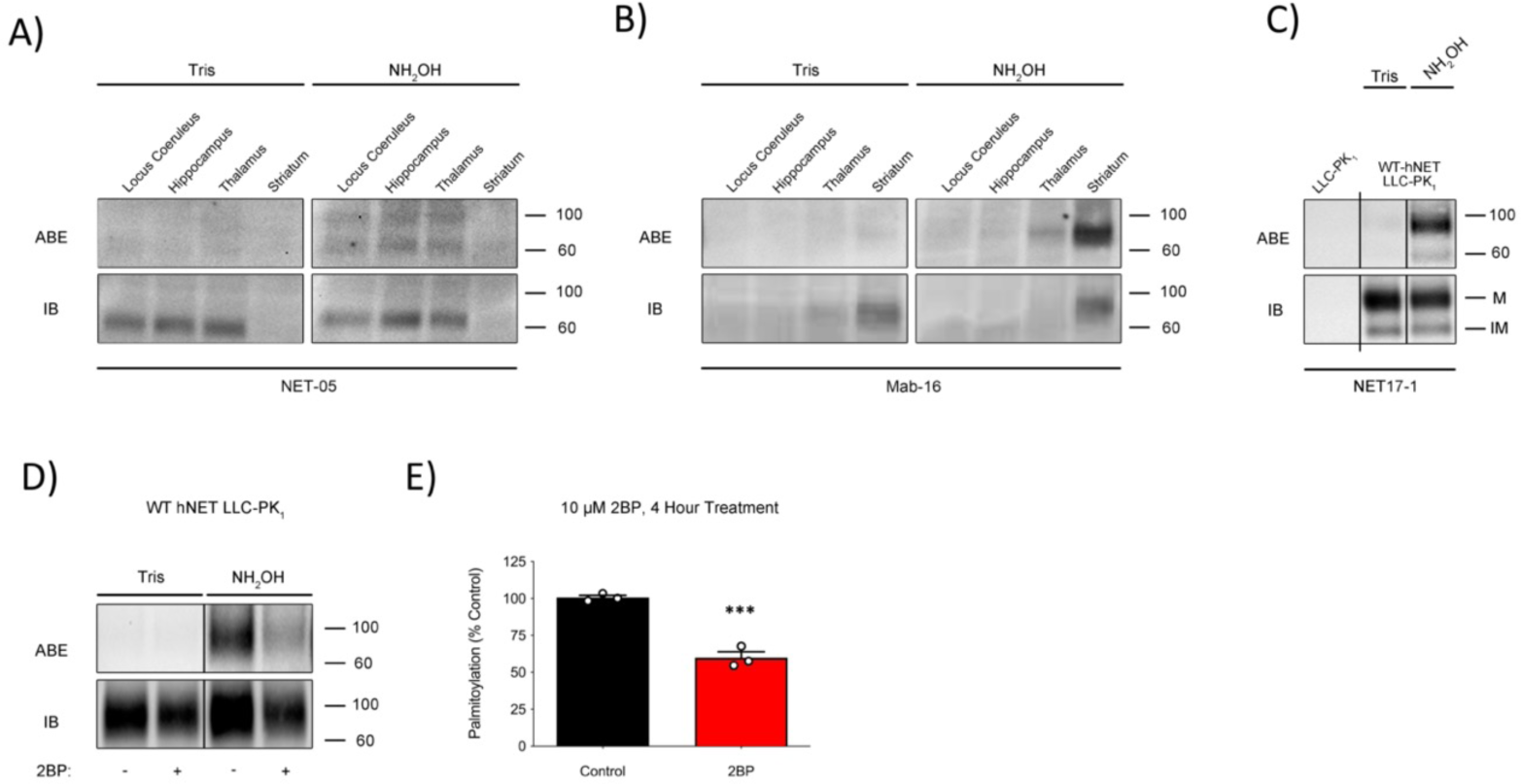
S-palmitoylation of rat and human NET and inhibition by 2BP. (***A, B***) Rat brain synaptosomes and **(*C)*** LLC-PK_1_ cells expressing WT hNET were assessed for palmitoylation (ABE) followed by SDS-PAGE and immunoblotting with either anti-mouse NET-05 or anti-human NET17-1 Ab, respectively. Samples were treated with Tris-HCl, pH 7.4 (control), or NH_2_OH to remove thioester-linked palmitate, followed by sulfhydryl-reactive biotinylation, NeutrAvidin extraction, and NET immunoblotting. Total NET protein (IB) in ABE samples taken just prior to NeutrAvidin extraction was also assessed via immunoblotting **(B)** Rat brain synaptosome immunoblots were subsequently stripped and re-probed with anti-rat DAT MAb-16 Ab. **(*D*)** LLC-PK_1_ cells expressing WT hNET were treated with vehicle or 10 µM 2BP for 4 h followed by assessment of palmitoylation (ABE) and total hNET levels (IB) in ABE samples. Extended black lines (|) on the top and bottom of panel C indicate two separate blots while black lines within the boundaries of the image of panels C and D indicate the removal of duplicate lanes or rearrangement of lane images from the same blot. **(*E*)** Quantification of NET palmitoylation normalized to total NET protein present in each sample (Mean ± SEM of 3 independent experiments (n=3) relative to control normalized to 100%; *** p < 0.001 versus Control by Student’s t-test).

To better understand the nature of the thioester modification, we treated our cells for 4 h with 10 µM 2-bromopalmitate (2BP) – an irreversible global inhibitor of the palmitoyl acyl-transferase family of enzymes (PATs, DHHCs). Immunoblotting revealed a strong 40 ± 4% reduction in hNET palmitoylation relative to control (*Fig. 1D and E*) (p<0.001 via Student’s t-test, n=3). Analysis of the total hNET in these immunoblots (i.e., Fig 4D, lower panel (IB); *data not shown*), revealed an ∼30% decrease in total NET expression under these conditions (p<0.01 via Student’s t-test, n=7). This suggested to us that a potential mechanism may exist where palmitoylation prevents the loss of total NET, either through opposing degradation or increasing biogenesis. Collectively these results suggest that rNET and hNET are palmitoyl-proteins, that hNET is sensitive to inhibition by 2BP, and that this modification modulates total hNET protein levels.

### Acute Inhibition of NET Palmitoylation Decreases Total NET, Surface Expression, and Transport Capacity

Following our observation that 4 h of 2BP treatment induced a loss in NET palmitoylation, we focused our efforts to determine the initial onset of this process. To characterize this timeframe, we treated cells expressing WT hNET with 7.5 µM 2BP and assessed palmitoylation over 120 min of incubation. ABE analysis of these samples revealed that palmitoylation of NET was reduced to 53.8 ± 17.7% of baseline within 90 min of 2BP treatment (*Fig. 2A and B*; p<0.05 via ANOVA with Tukey post hoc test, n=3). This decrease remained through 120 min at 58.5 ± 10.4% of basal conditions (*Fig. 2A and B*; p<0.05 via ANOVA with Tukey post hoc test, n=4). Analysis of the post-ABE total NET protein revealed a reduction in total NET beginning at 120 min of 2BP treatment, with 75.4 ± 4.4% of total NET protein remaining (*Fig. 2A and C*; p<0.05 via ANOVA with Tukey post hoc test, n=4). The observation that 90 min of 2BP treatment decreased NET palmitoylation which continued through 120 min and was accompanied by the loss of total NET protein after 120 min suggested that a relationship exists between losses in palmitoylation and the stability of NET expression.

**FIGURE 2.**
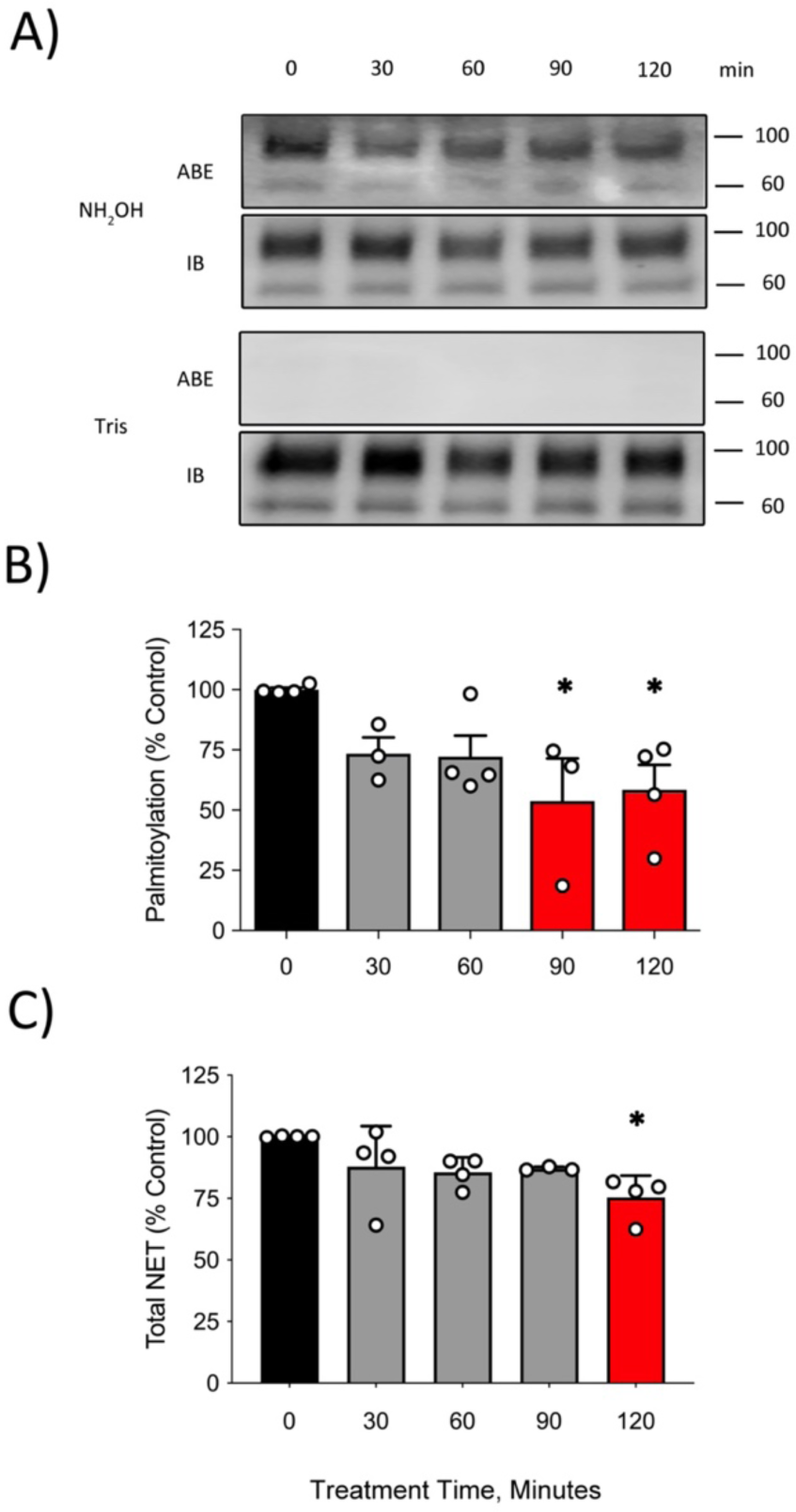
Acute inhibition of NET palmitoylation by 2BP reduces total NET protein. LLC-PK_1_ cells expressing WT hNET were treated with 7.5 µM 2BP for 0, 30, 60, 90 and 120 min followed by assessment of palmitoylation levels by ABE. Total NET protein (IB) in ABE samples taken just prior to NeutrAvidin extraction was also assessed via immunoblotting **(*A)*** Western blot representative of three or more independent ABE experiments for each time point after treatment with 7.5 µM 2BP with corresponding immunoblot for total NET protein levels (IB) used to normalize palmitoylated NET levels. **(*B*)** Quantification of NET palmitoylation (Mean ± SEM of three or more independent experiments performed relative to control (Treatment Time, 0) normalized to 100%.). * p < 0.05 versus time point 0 (one-way ANOVA with Tukey post hoc test). **(*C*)** Quantification of post-ABE NET total expression blots (Mean ± SEM of at least 3 independent experiments relative to control (Treatment Time, 0 min) normalized to 100%). * p < 0.05 versus time point 0 (one-way ANOVA with Tukey post hoc test).

Our next step was to determine if the loss in total NET protein would correlate with changes in NET surface expression and transport capacity. To test for changes in NET trafficking, we analyzed NET surface expression *via* cell surface biotinylation and DA transport after challenge with 7.5 µM 2BP. Selection of DA as the substrate for uptake was based on data describing NET as the predominant reuptake mechanism for DA in the prefrontal cortex (35), its robust affinity for DA, and our ease of accessibility as we routinely use [^3^H]DA in our DAT studies. Demonstrated in Figure 3A and B, 120 min of 2BP treatment decreased NET surface expression to 77.0 ± 4.2% of vehicle control that mirrored the time course for decreased palmitoylation and total NET protein (p<0.05 via ANOVA with Tukey post hoc test, n=3-6). DA uptake analysis revealed a nearly identical trend in loss of transport capacity as seen with NET surface and total cellular expression (*Fig. 3C*; 64.9 ± 7.9%, p<0.05 via ANOVA with Dunnett post hoc test, n=4). These data suggest that palmitoylation is responsible for acute maintenance of NET surface expression, total protein, and transport capacity.

**FIGURE 3.**
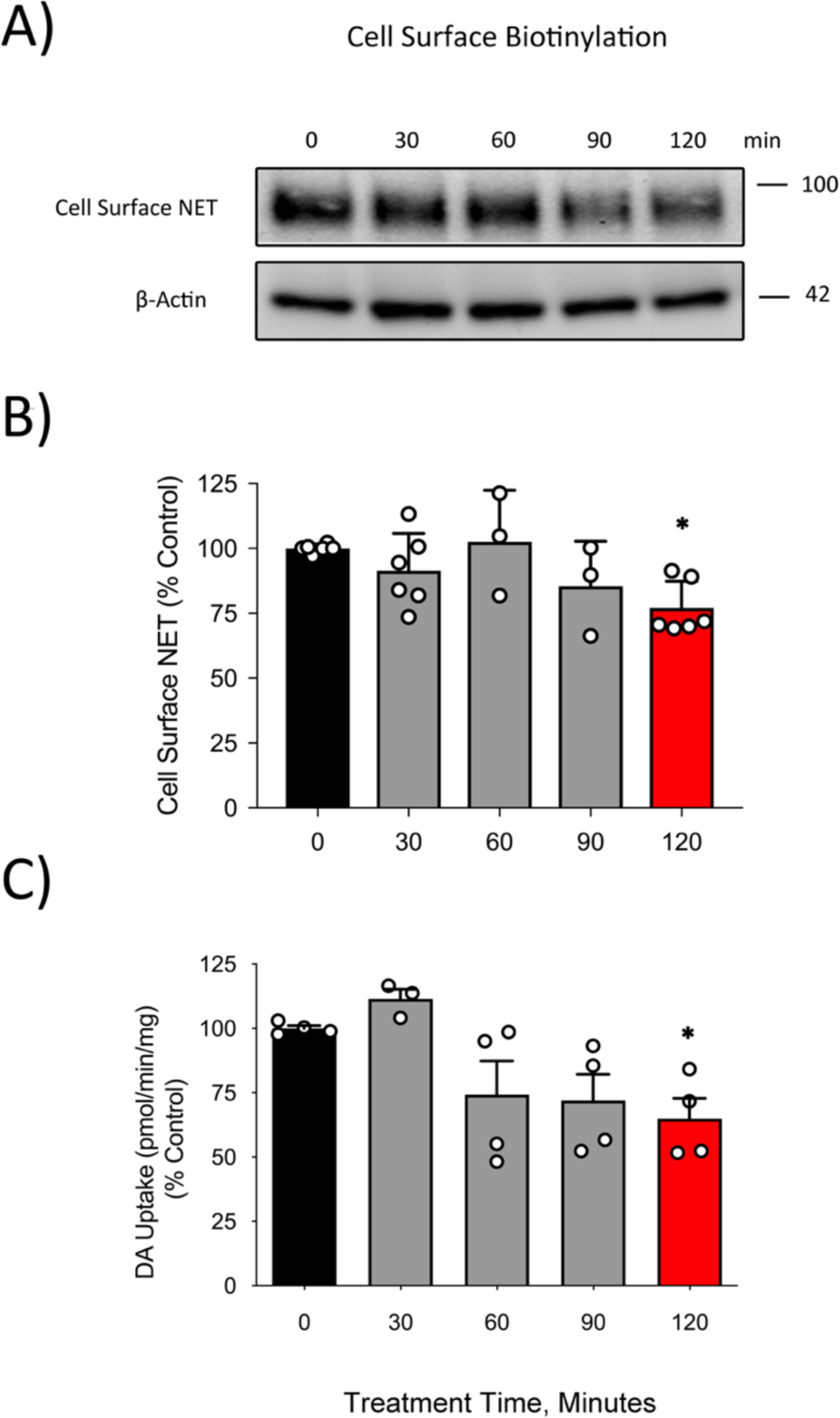
2BP decreases NET surface expression and transport capacity. LLC-PK_1_ cells expressing WT hNET were treated with 7.5 µM 2BP for the indicated times followed by assessment of hNET cell surface expression and DA uptake. **(*A*)** A representative immunoblot of hNET in biotinylated fractions eluted from NeutrAvidin resin and β-Actin levels in equal protein loads of cell lysates from at least 3 independent experiments (n = 3-6) for each time point. **(*B*)** Quantification of cell surface hNET blots normalized to β-Actin expression in lysates from cell surface biotinylation. (Mean ± SEM of at least 3 independent experiments relative to control (Treatment Time, 0 min) normalized to 100%; * p < 0.05 versus time point 0 (one-way ANOVA with Tukey post hoc test). **(*C*)** Quantification of DA uptake by NET following treatment times with 2BP (Mean ± SEM of 4 independent experiments relative to control (Treatment Time, 0 min) normalized to 100%; * p < 0.05 versus time point 0 min (one-way ANOVA with Dunnett post hoc test).

Collectively, these results suggest that a relationship exists between palmitoylation, trafficking, and degradation and/or biogenesis of NET protein. With such an acute impact, it’s possible that when NET palmitoylation is inhibited by 2BP, NET is prevented from remaining at the cellular surface either by internalization or decreased surface recruitment. This process would result in retention of endosomal NET with subsequent sorting to late endosomes and lysosomal degradation, reducing its ability to reuptake extracellular catecholamines.

### 2BP Inhibition of Palmitoylation Does Not Impact NET Transport Kinetics

Our results suggested that palmitoylation regulates cell surface trafficking, total expression, and transport capacity of NET (*Fig. 2 and 3*). Thus, we wanted to determine if the decreased NET transport capacity at 120 min was due to changes in NET transport kinetics as we have described for DAT and SERT (26,29,30) or exclusively the result of decreased NET surface density. To study this, we utilized DA uptake saturation analysis and calculated K_m_ and V_max_ normalized to total cellular protein (pmol/min/mg protein) and parallel cell surface biotinylation analysis to determine NET surface levels (*Fig. 4*) and normalize DA uptake. This method and normalization allows us to distinguish between kinetic or trafficking effects in response to 2BP inhibition of NET palmitoylation as we have reported previously for DAT and SERT (26,29,30). LLC-PK_1_ cells expressing WT hNET treated with vehicle (DMSO) or 7.5 µM 2BP for 30 or 120 min showed no change in surfaced normalized V_max_ or in K_m,DA_ under these condition (*Fig. 4A-D;* p >0.05, ANOVA with Tukey post hoc test) demonstrating no kinetic modulation of NET function by palmitoylation.

**FIGURE 4.**
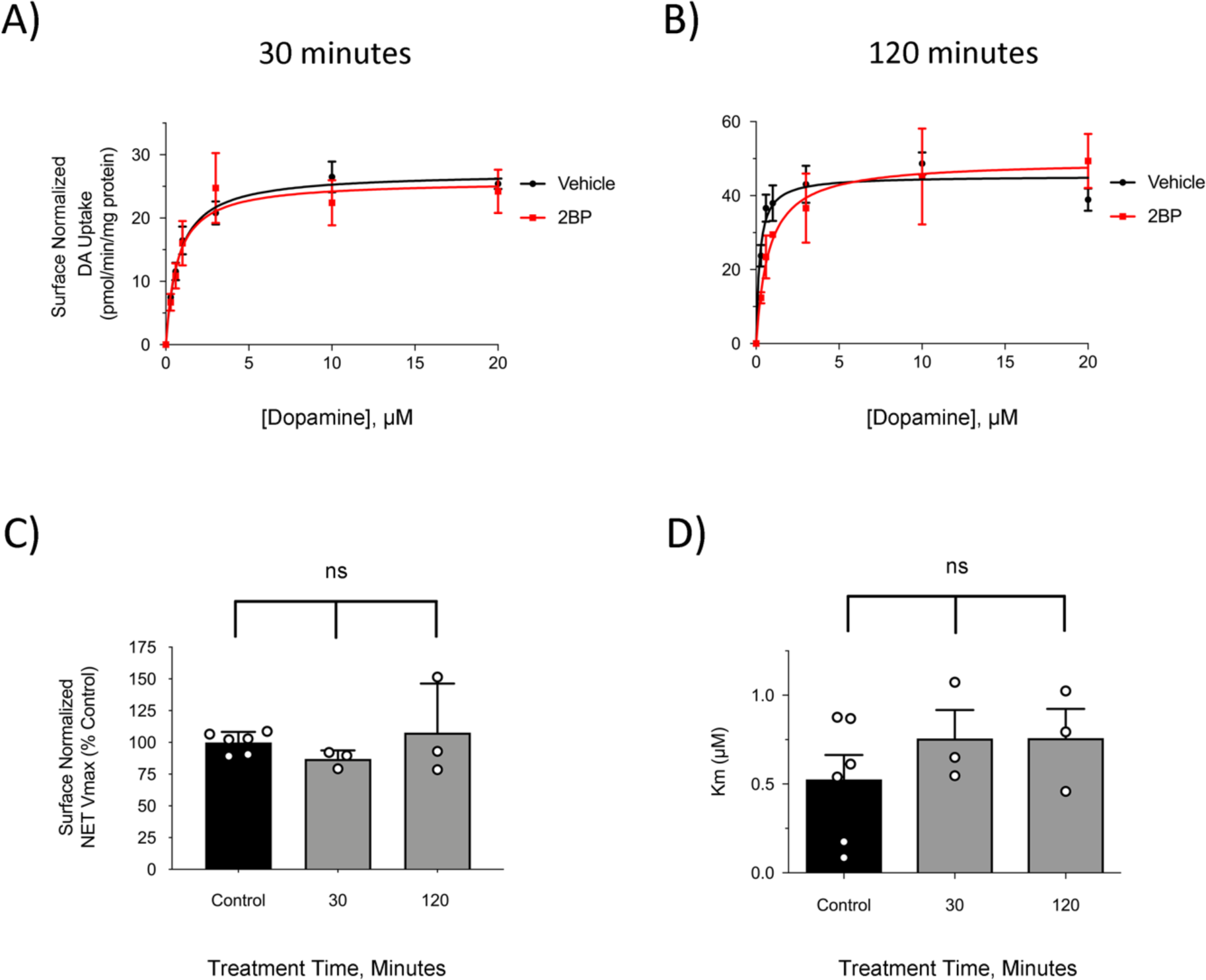
NET transport kinetics are unchanged with differential 2BP time treatments. LLCPK_1_ cells expressing WT hNET were treated with vehicle or 7.5 µM 2BP for 30 min **(*A*)** or 120 min **(*B*)** were subjected to DA transport saturation analysis presented as means ± SEM of three independent experiments performed in triplicate. ***(C)*** Quantification of surface normalized V_max_ (Mean ± SEM of at least 3 independent experiments relative to control normalized to 100%.). ***(D*)** Quantification of K_m, DA_ values (Mean ± SEM of at least 3 independent experiments relative to control).

Treatment of WT hNET with 7.5 µM 2BP decreased palmitoylation within 90 min that continued through 120 min (*Fig. 2A, B*). However, we did not detect any statistically significant changes in total NET, surface NET, or NET-mediated DA uptake until 120 min of 2BP treatment. NET palmitoylation appears to acutely regulate catecholamine uptake through trafficking-dependent changes while palmitoylation of DAT and SERT functions to directly modulate the transporters’ kinetics within 30 min of 2BP treatment (29,30). Ultimately, these data highlight that NET expression is exceptionally sensitive to acute changes in palmitoylation that control its trafficking and stability, underlining striking distinctions that exist between NET, DAT, and SERT regulation by palmitoylation.

### 2BP-Mediated Inhibition of NET Protein Expression

Following our observation that acute inhibition of NET palmitoylation with 2BP led to a rapid decline in total NET protein, we expanded our 2BP treatment times and concentrations. We have previously identified that chronic inhibition of DAT palmitoylation *via* 2BP or site-directed mutagenesis of palmitoylated-cysteines diminishes total DAT protein expression that is often accompanied with DAT degradation fragments (26). Additionally, we have demonstrated that driving palmitoylation *via* overexpression of DHHC enzymes leads to increased total DAT expression (29), supporting the notion that palmitoylation controls long-term regulation of DAT by opposing lysosomal turnover and potentially facilitating biogenesis.

In this regard, we extended our studies to include times ranging from 1-18 h at a constant concentration of 7.5 µM 2BP. Following this set of experiments, we challenged NET with progressively increasing 2BP concentrations ranging from 1-50 µM with 18 h of exposure. These studies were intended to deepen our understanding of the impact that inhibition of palmitoylation has on NET protein expression and trafficking. Within 2 h of 2BP-mediated inhibition of NET palmitoylation, total NET protein was significantly decreased to 84.2 ± 4.4% of control (Fig. 5*B*, p<0.05 via ANOVA with Tukey post hoc test). This trend continued in a time-dependent manner with total NET protein dropping to 26.7 ± 2% at 18 h (Fig. 5*B*, p<0.0001 via ANOVA with Tukey post hoc test). The irreversible nature of the inhibitor 2BP is well demonstrated with the time-dependent decrease in NET protein over the 18 h period, with no plateau effect or recovery of NET protein within the treatment time. Notably, there was no change in quantity or integrity of β-actin for each time point, suggesting that 2BP’s impact on NET was not in response to generalized degradative processes (*data not shown*). These data suggest that palmitoylation is critical for maintenance of NET stability through stabilizing biogenesis and/or opposing degradation.

**FIGURE 5.**
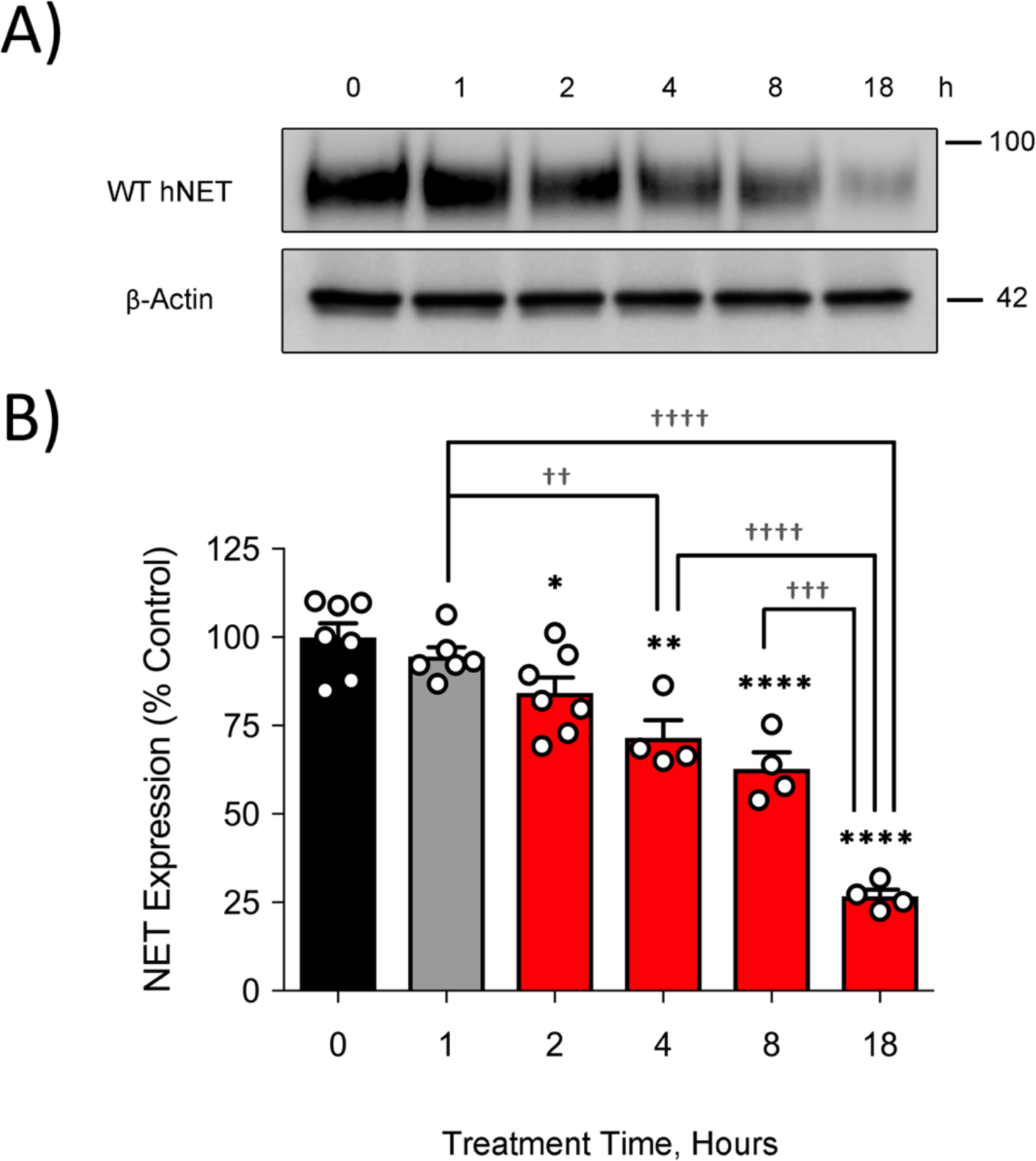
Time-dependent loss of total NET protein induced by 2BP. LLC-PK_1_ cells expressing hNET were treated with 7.5 µM 2BP for 0, 1, 2, 4, 8, and 18 hours followed by membrane preparation and determination of total NET protein levels by immunoblotting equal membrane protein loads (IB) **(*A*)** Western blot representative of at least four (n = 4-7) independent experiments of NET following treatment with 2BP. Representative blot of β-actin levels from the same cells in equal protein loads of the membrane fraction are also demonstrated. **(*B*)** Quantification of hNET blots (Mean ± SEM of 4-7 independent experiments relative to control (Treatment Time, 0 h) normalized to 100%. * p < 0.05, ** p < 0.01, **** p < 0.0001 versus control; ^†^ p < 0.05, ^†††^ p < 0.001, ^††††^ p < 0.0001 between indicated concentrations (one-way ANOVA with Tukey post hoc test).

We next wanted to determine the maximal extent to which total NET expression could be reduced. Theoretically, since 2BP is an irreversible inhibitor of all DHHC family members, there should be a concentration-dependent decrease in total NET expression that would resemble our time-dependent findings in Figure 5. To examine this, we challenged cells expressing WT hNET with 2BP at progressively increasing concentrations of 2BP between 1-50 µM for 18 h. Figure 6A outlines a dramatic loss in total NET protein when treated with as little as 1 µM 2BP for 18 h (51.9 ± 3.3% of Control, p<0.0001 *via* ANOVA with Tukey post hoc test) that continued in a concentration-dependent manner through 50 µM 2BP (22.8 ± 1.9% of Control, p<0.0001 *via* ANOVA with Tukey post hoc test). Importantly, no change in β-actin expression was observed with all 2BP concentrations used suggesting that high concentrations of 2BP do not trigger significant cellular cytotoxicity in LLC-PK_1_ cells consistent with our previous work using this cell line (26). Overall, these data demonstrate that acute inhibition of palmitoylation and decreased total NET protein continues through 18 h of treatment and nearly abolishes the total level of NET protein after treatment with a high concentration of 2BP. These data also suggest that palmitoylation is essential in maintenance of total NET protein, either through facilitating proper biogenesis and/or opposing lysosomal degradation.

**FIGURE 6.**
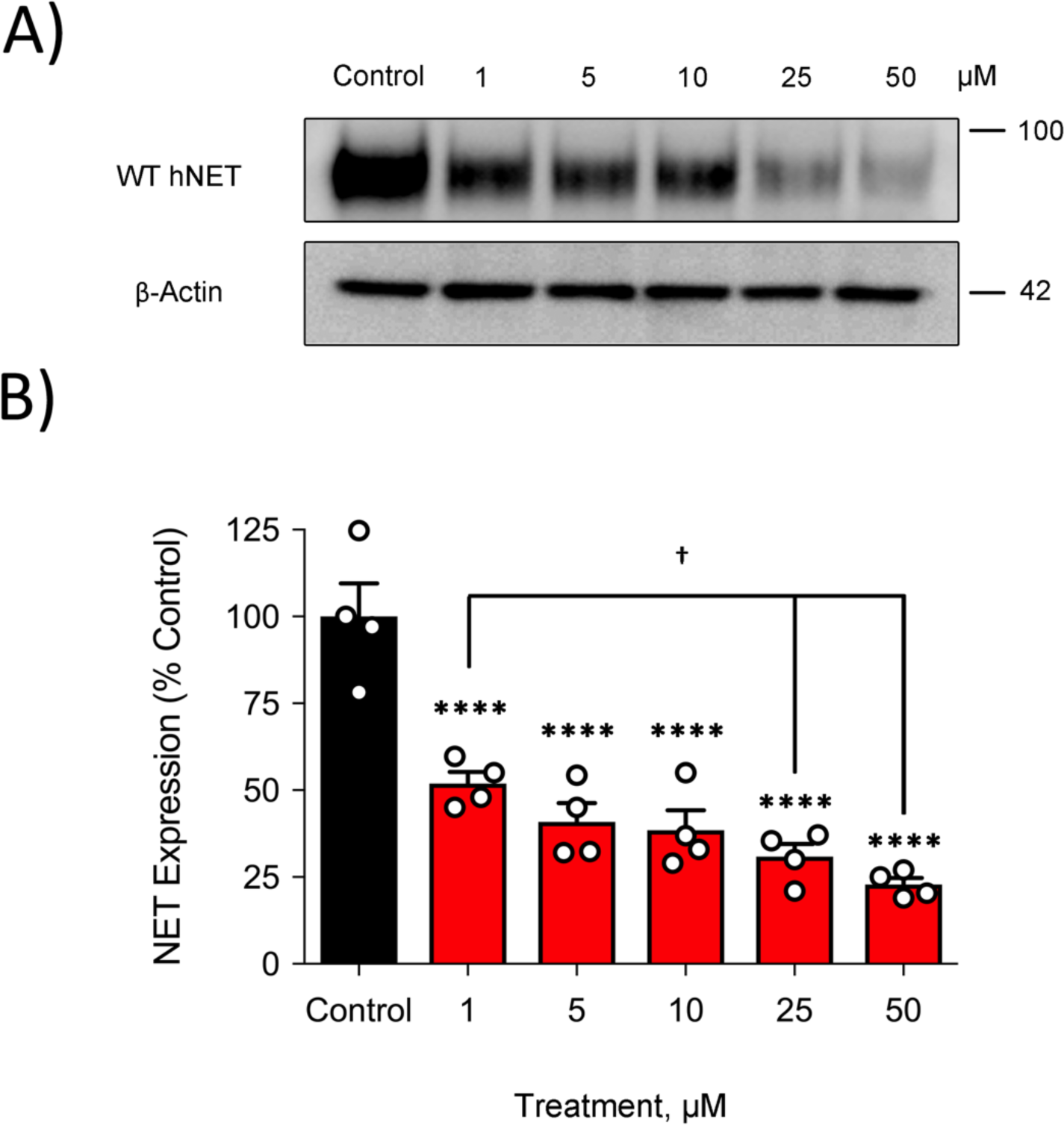
Concentration-dependent loss of total NET protein induced by 2BP. LLC-PK_1_ cells expressing hNET were treated with 0, 1, 5, 10, 25, and 50 µM 2BP for 18 hours followed by membrane preparation and determination of total NET protein levels by immunoblotting equal membrane protein loads (IB) **(*A*)** Western blot representative of four independent hNET experiments for each concentration of 2BP. A representative blot of β-actin levels from these same cells with equal protein loads of the membrane fraction is also demonstrated. **(*B*)** Quantification of hNET blots (Mean ± SEM of at least 4 independent experiments relative to control (Treatment Concentration, 0 µM) normalized to 100%. **** p < 0.0001 versus control; ^†^ p < 0.05 (one-way ANOVA with Fisher-LSD post hoc test).

### DHHC1 Overexpression Stimulates NET Palmitoylation

Similar to our previous studies examining DAT palmitoylation, we assessed DHHC-specific NET palmitoylation utilizing a DHHC overexpression approach (29). Because specific inhibitors are not available for each of the 23 DHHC enzymes (36,37) we chose to circumvent this problem by employing a co-expression strategy using DNA plasmids coding for HA-tagged mouse DHHC proteins (38). LLC-PK_1_ cells stably expressing WT hNET were transfected with DHHC enzymes that that were previously shown to enhanced DAT palmitoylation, kinetics, and total DAT levels (DHHC 2, 3, 8, 15, and 17), including an additional DHHC enzyme, DHHC1 (29). In unpublished data, we have found that DHHC1 promotes SERT palmitoylation, and was thus included in the current studies’ experimental design. Additionally, DHHC1 is an ER bound enzyme, suggesting its importance in protein biogenesis and maturation (39). This was considered a potential mechanism, and tested accordingly, to help explain our current results that suggest palmitoylation may be necessary for NET biogenesis and stability.

To analyze NET palmitoylation, cells underwent transient transfection with the indicated DHHC cDNA plasmid and 24 h post-transfection, palmitoylation was assessed *via* ABE and normalized to total NET protein present. A representative western blot shown in Figure 7A demonstrates an enhancement in NET palmitoylation by DHHC1 (157.3 ± 9.9% of control *via* one way ANOVA with Tukey post hoc test, n=4, p<0.05). Although not statistically significant, expression of DHHC2 suggested a trend towards an increase in NET palmitoylation (119.2 ± 8.9% of control). The other DHHCs tested (DHHC3, 8, 15, 17, 20, 21, and 22) had no statistical impact on NET palmitoylation when compared to the control sample. DHHC expression was confirmed by immunoblotting with anti-HA monoclonal antibody to detect the HA-tagged DHHCs (*Fig. 7B, bottom panel*). The histogram (*Fig. 7C*) outlines the collective findings of these experiments. Although this list is not all inclusive for the DHHC family (37,39), and DHHC1 was not tested in our previous studies with DAT (29), we find it interesting that the DHHC members presently investigated appear to have more limited selectivity for driving NET palmitoylation than DAT, where co-expression of DHHCs 2, 3, 8, 15 and 17 were found to stimulate DAT palmitoylation (29).

**FIGURE 7.**
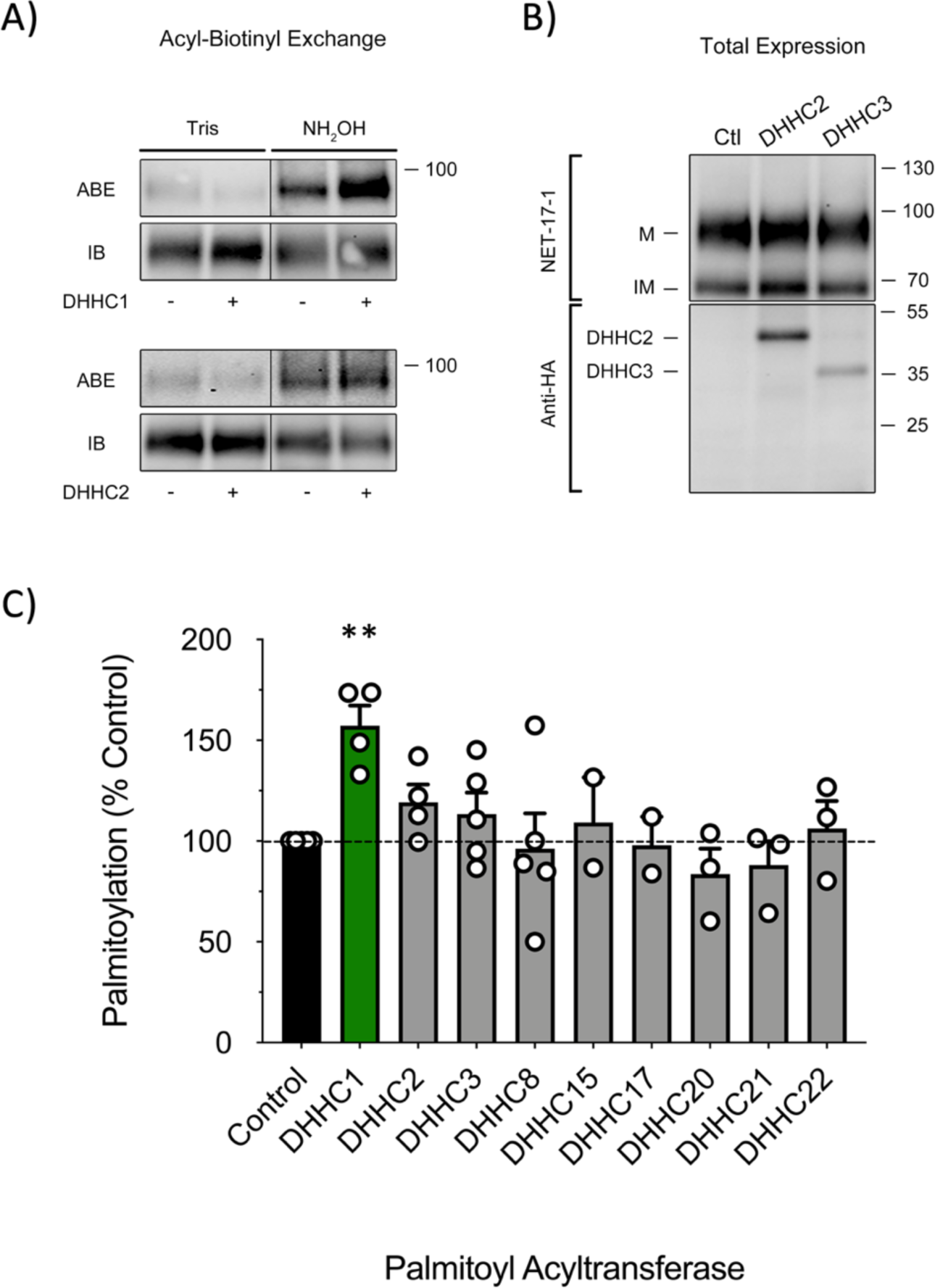
NET palmitoylation levels following differential DHHC expression. LLC-PK_1_ cells expressing hNET were transfected with the indicated DHHC plasmids and assessed for NET palmitoylation by ABE 24 h post transfection. **(*A*)** Immunoblots show representative ABE (palmitoylated) and total hNET in samples (IB) of at least four independent experiments for the indicated DHHC transfection. Black lines (|) within the boundaries of the images of panel A indicate the removal of duplicate lanes or rearrangement of lane images from the same blot. **(*B*)** Western blot representative of WT hNET (M, mature; IM, immature) following DHHC2 and 3 transfection (upper panel – blotted with Anti-hNET (NET-17-1)) and western blot representative of HA-tagged-DHHC2 and 3 expression following transfection (lower panel – blotted with Anti-HA Ab) **(*C*)** Quantification of NET palmitoylation (Mean ± SEM of at least 2 (n=2-5) independent experiments relative to control normalized to 100%.) ** < 0.01 versus Control (one-way ANVOA with Fisher-LSD post hoc test).

### DHHC1 Co-expression Increases Membrane-associated A457P Fragmentation

Our lab has previously demonstrated that driving DAT palmitoylation by DHHC co-expression results in increased DAT cellular expression (29). Because our current study demonstrates DHHC1 co-expression enhances NET palmitoylation (Fig. 7) and inhibition of palmitoylation with 2BP rapidly and dramatically decreases total NET protein (*Fig. 2, 5, and 6*), we thought that DHHC1 co-expression may result in enhanced total cellular NET protein. To test this, we transfected LLC-PK_1_ cells stably expressing WT hNET with DHHC1 and assessed NET protein levels by immunoblotting equal amounts of membrane protein which revealed an increase in total NET protein of 131.2 ± 8.7% compared to the WT control group (*Fig. 8A*, p<0.05 *via* one way ANOVA with Tukey post hoc test, n=3).

**FIGURE 8.**
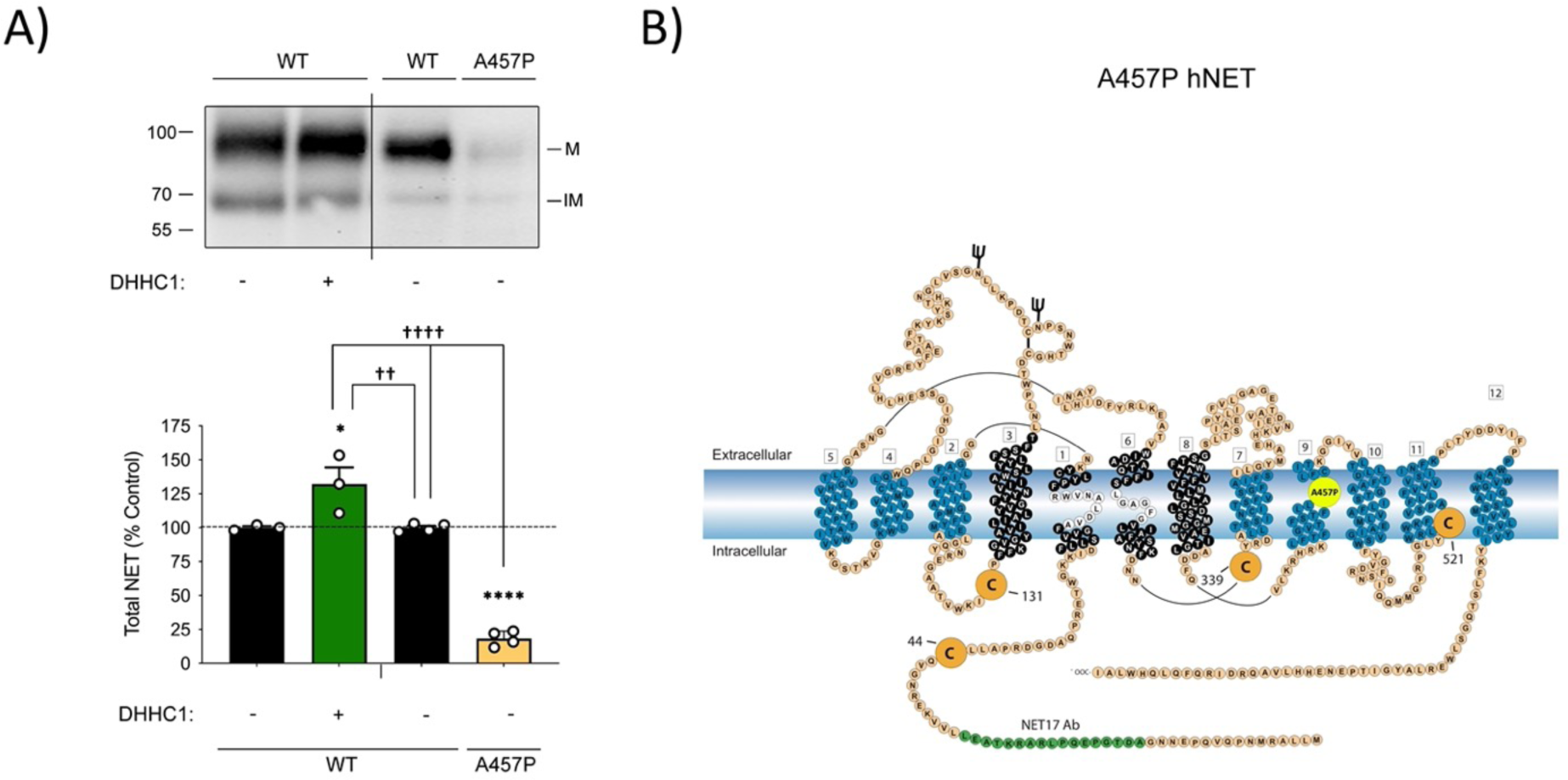
Total protein levels of A457P and WT hNET. (***A***) Equal protein loads of membranes isolated from LLC-PK_1_ cells expressing WT hNET or A457P hNET with (+) and without (-) transient transfection with DHHC1 were resolved by SDS-PAGE and immunoblotted with anti-NET Ab (NET-17-1; M, mature; IM, immature). Black lines (|) extended beyond the boundaries of the images of panel A indicate two separate blots. The histogram shows quantification of total NET protein under the indicated type and/or condition (Mean ± SEM of at least three independent experiments (n=3-4) performed relative to WT control [left-most WT, (-)] normalized to 100%). * p < 0.05, **** p <0.0001 vs WT control; ^††^ p < 0.01, ^††††^ p < 0.0001 vs WT control (one-way ANOVA with Tukey post hoc test). **(*B*)** hNET is composed of twelve transmembrane spanning domains that are connected by alternating intracellular and extracellular loops. Large orange circles denote potential intra-cellular cysteine sites for palmitoylation. C44 is located distally to the N-terminally located NET-17-1 epitope. A457P substitution is identified by the large yellow circle within the 9^th^ transmembrane domain.

Additionally, previous studies have implicated NET via the coding variant A457P in the pathogenesis of POTS (17). In those studies, the dramatic decrease in total A457P expression was suggested to occur from a disturbance in NET biogenesis at the level of ER-Golgi processing. This could potentially result in decreased synthesis, glycosylation, and recruitment of NET to the cell surface. Following our NET palmitoylation studies, we recognized there were striking similarities in our findings and the findings of prior research (17,18,40) regarding the expression of NET. In accordance with our current findings, where inhibition of NET palmitoylation resulted in a dramatic and rapid loss of NET protein (*Fig. 5 and 6*), we hypothesized that palmitoylation of A457P NET at the ER-Golgi level may play a role in the maturation and expression of NET. The low amounts of A457P expression prevented us from reliably determining palmitoylation levels in the A457P variant so we attempted to drive palmitoylation of the variant by DHHC1 co-expression which we showed enhanced WT NET palmitoylation and expression levels (Figs 7 and 8A). To test this, we established cells stably expressing A457P NET and assessed total NET protein levels. Equal amounts of membrane protein were resolved by SDS-PAGE and immunoblotted for NET using anti-NET-17-1 Ab. The results of this experiment confirmed that A457P substitution results in a dramatic reduction in total expression (*Fig. 8A*, 18.4 ± 2.8%, p<0.0001 *via* one way ANOVA with Tukey post hoc test, n=4) confirming previous reports (17,18). A topographic representation of hNET is shown in Figure 8B, were the A457P substitution is found in the 9^th^ transmembrane domain and facilitates a massive reduction in total and cell surface transporter expression. To test our hypothesis that driving palmitoylation with DHHC1 co-expression would recover expression of mature NET, we co-expressed DHHC1 in cells stably expressing A457P NET. Unexpectedly, co-expression of DHHC1 with A457P NET did not increase the mature glycosylated fraction of A457P NET but was accompanied by the presence of immunoreactive N-terminal NET fragments (*Fig. 9A*). Quantification of all immunoreactive bands in the A457P blots in the presence or absence of DHHC1 expression revealed a DHHC1-dependent increase in A457P NET immunoreactivity by 25 ± 2.1% compared to A457P control (*Fig 9A*, p<0.00001 *via* ANOVA with Tukey post hoc test). It is noteworthy to mention that equal amounts of protein from membrane preparations, and not whole cell lysates, were resolved by SDS-PAGE suggesting the NET fragments are membrane localized.

**FIGURE 9.**
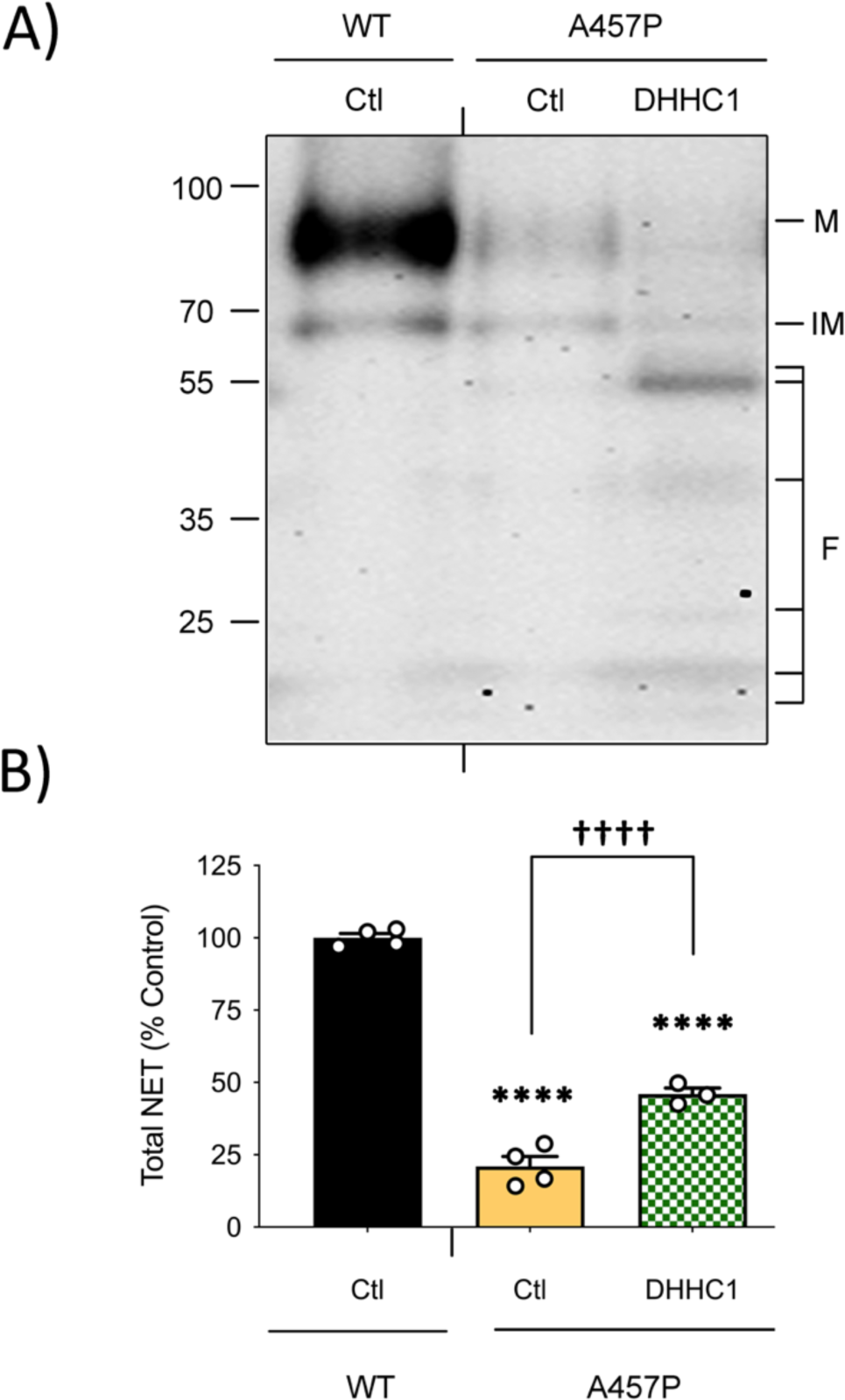
DHHC1 expression enhances A457P fragmentation. Equal protein loads of membranes isolated from LLC-PK_1_ cells expressing WT hNET (Ctl) or A457P hNET without (Ctl) or with DHHC1 transfection (24h) were resolved by SDS-PAGE and immunoblotted with anti-NET Ab (NET-17-1) **(*A*)** Western blot representative of three or more independent experiments for WT and A457P hNET total protein (M, mature; IM, immature; F, fragments). **(*B*)** Quantification of the entire column for total NET protein (Mean ± SEM of three or more independent experiments performed relative to WT control (Ctl) normalized to 100%.) **** p<0.0001 versus WT Ctl, ^††††^ p<0.0001 versus A457P Ctl (one-way ANOVA with Tukey post hoc test).

## DISCUSSION

NET has been characterized as a target for many post-translational modifications including phosphorylation, glycosylation, and ubiquitylation (41). Phosphorylation of NET by PKC has been demonstrated to occur at T258 and S259, and functions to decrease NET surface expression and transporter V_max_, without changing the K_m NE_ (42,43). In the presence of cocaine, NET is up-regulated through a p38 MAPK-dependent process, where phosphorylation of T30 inhibits transporter endocytosis and increases NET surface density and activity (44). Through N-glycosylation, NET is directed to the cell surface to perform its physiologic function (40). When glycosylation is inhibited via mutation or enzymatically cleaved, the resultant unglycosylated NET remains in intracellular compartments (40). Likewise, unglycosylated NET protein was reduced ∼50% when compared with glycosylated NET, suggesting that glycosylation is important in maintaining NET stability and/or preventing degradation (40). These modifications work collectively to modulate NET availability and activity in correspondence with physiologic requirements, providing a mechanism to maintain catecholamine homeostasis in a consistently changing physiologic landscape. Our previous studies on the post translational modification S-palmitoylation have revealed an additional layer of regulation for DAT (26) and SERT (30). These previous studies have outlined palmitoylation as a major player in acute and chronic modulation of both transporter’s kinetic activity, trafficking, and total transporter protein levels. We expanded our scope here to include NET, as all three monoamine transporters exhibit similar membrane topography, amino-acid core sequences, and execute similar physiologic functions.

In this study, we reveal that palmitoylation is an essential component in the dynamic mechanisms of NET regulation. Using the palmitoylation inhibitor, 2BP, we outline several functional consequences of palmitoylation that have temporal and concentration-based differences. In acute time frames, NET palmitoylation was found to be inhibited after 90 min of 2BP treatment that continued through 120 min. In contrast to DAT and SERT, we discovered that acute inhibition of palmitoylation promoted a loss in total NET protein within 120 min of treatment. In other words, compared with NET, DAT and SERT are less sensitive to changes in total transporter levels when palmitoylation is inhibited acutely, requiring extended (∼18 h) inhibition of palmitoylation to result in decreased transporter protein levels (26,30). To date, palmitoylation has not been observed to modulate DAT trafficking properties (26,29). However, when DAT is challenged with 2BP in chronic conditions (18 h), we observed a loss in total DAT protein that is accompanied with DAT degradation fragments (26) suggesting that DAT trafficking is likely influenced by palmitoylation. With SERT, we have observed that inhibition of palmitoylation with 2BP for 3 h triggers a decrease in surface SERT that ultimately leads to decreased total SERT protein levels with 18 h of treatment (30). Acutely, inhibition of palmitoylation did reduce DAT and SERT V_max_ independent of changes in cell surface expression (26,29), indicating kinetic effects that are not seen with NET in this study. Overall, these results lead to the interesting conclusion that palmitoylation regulates NET, DAT, and SERT with different outcomes, with palmitoylation acutely controlling NET protein stability and trafficking, DAT kinetics, and SERT trafficking and kinetics (26,29,30,45). This difference is likely the result of differences in the number of palmitoylation sites and their location in each transporter, the DHHC enzyme that catalyzes palmitoylation at each site, and the subcellular location where the modification occurs.

Consistent with palmitoylation and total protein levels, we observed a loss in NET surface expression and transport capacity after 120 min of 2BP exposure. These data suggest that NET trafficking is largely dependent on its palmitoylation status, potentially protecting NET against degradative mechanisms and/or promoting surface recruitment. This seems possible, as our data on DHHC co-expression with NET suggests that DHHC1 likely catalyzes the palmitoylation of NET, increasing total NET expression. Interestingly, DHHC1, an integral membrane protein, has been determined to be localized to the ER membrane (39,46), suggesting that DHHC1 may enhance NET palmitoylation in a co-or post-translational process that facilitates appropriate biogenesis and processing of the transporter. This is not out of the realm of possibility, as other transporters, channels, and surface proteins exhibit a similar method of post-translational development (27,28,31). The cystic fibrosis transmembrane regulator (CFTR) is a palmitoyl-protein that is dependent on DHHC7-mediated palmitoylation in the ER compartment, where defects in CFTR biogenesis are central to the pathogenesis of cystic fibrosis. This study outlined that 2BP decreases the total expression of both WT and low-temperature corrected F508del CFTR – decreasing post ER-processing, CFTR surface expression, and ion channel activity (31). Even more so, the investigators found that this process could be rescued through DHHC7 overexpression, increasing total CFTR expression for both WT and F508del, and sequestration of CFTR in post-ER compartments (31). In this model, the majority of palmitoyl-CFTR was found to be sequestered in the Golgi, suggesting that another post-translational event was necessary for subsequent maturation and glycan attachment. These results are highly suggestive that at least one function of DHHCs is to direct transmembrane protein processing and biogenesis.

The POTS NET coding variant, A457P, drew our attention after we determined NET protein levels are extremely sensitive to 2BP-mediated inhibition of palmitoylation. The characteristics of A457P and our NET palmitoylation studies highlight several key similarities. For example, 2BP induces a rapid decrease in cell surface and total NET expression that was consistent with palmitoylation levels, ultimately decreasing NET transport capacity. However, our saturation analysis indicated no change in V_max_ or K_m DA_ of the transporter after inhibition of palmitoylation with 2BP. These data are suggestive that inhibition of NET palmitoylation decreases the density of NET at the cellular surface, either by internalization or inhibition of cell surface recruitment. Regardless, this process results in a reduction in transport capacity without accompanying direct kinetic changes, implying that palmitoylation controls NET transport largely through a trafficking-based mechanism. A457P NET has demonstrated similar properties, exhibiting a large decrease in transport, surface, and total expression without changes in ligand binding properties (17). The investigators of this study hypothesized that NET processing was disturbed at the ER/Golgi level, because immature NET was seemingly unaffected in density with a corresponding decrease in mature NET protein levels (17) which is consistent with our current results (Fig. 8A). Although the maturation of transmembrane proteins is believed to involve N-glycosylation, these results suggested to us that palmitoylation of NET or ancillary proteins in the ER may impact NET’s maturation process. Consistent with previous studies, we demonstrated that A457P expression was reduced to ∼18% of WT levels (17,18). ER localized DHHC1 co-expression with A457P did not increase the mature form of total NET protein, but generated a striking increase in immuno-reactive NET N-terminal fragments that were localized to the membrane fraction of our cellular preparations. Even more interestingly, the NET-17-1 epitope is located proximal (N-terminal) to a likely palmitoylation site, C44, in the cytoplasmically facing N-terminal tail (see Fig. 8B) suggesting that palmitoylation may be anchoring these immunoreactive fragments to the various lipid membrane fractions of the cell. However, it is important to mention again that we cannot conclude with certainty that palmitoylation of A457P is decreased relative to the WT protein nor increases with DHHC1 co-expression, and thus responsible for this outcome, as total A457P levels are very low making ABE-mediated analysis of palmitoylation very difficult and unreliable. An alternative design to address this issue in future experiments includes mass-spectrometry based analysis of palmitoylated fractions of A457P under native conditions or the influence of DHHC1 co-expression. Using this approach would reveal specific protein sequence of the fragments and the fraction of palmitoylated NET under each experimental condition.

It seems possible that enhancing NET palmitoylation via ER-resident DHHC1 action may be involved in the physiologic sequence for biogenesis of NET, and that A457P disrupts this process *via* a conformational change. Because of the increased A457P NET fragmentation observed with DHHC1 co-expression, it is possible that driving palmitoylation with DHHC1 may transiently stabilize immature A457P processing through the ER-Golgi, but ultimately end with its degradation in the lysosome prior to reaching the cellular surface (*Fig. 10*). The mechanism for this, however, is unclear and could be multifaceted. One possibility is that NET glycosylation may be a prerequisite for palmitoylation in the ER. In this example, impaired glycosylation will decrease NET palmitoylation as a result of improper folding and subsequent steric hindrance. This would result in impaired formation of mature NET *via* glycosylation, and subsequent stability and surface trafficking from palmitoylation, consistent with previous studies and our current findings (17,18,40,47). This is supported by evidence that palmitoylation sorts membrane proteins for anterograde transport and trafficking from the Golgi (27). Another possibility is that palmitoylation may be a prerequisite for NET to undergo N-glycosylation, thereby promoting maturation and total cellular expression. This mechanism could be altered with the A457P variant, resulting in a strong reduction in its ability to be a target for palmitoylation, thus shunting the immature and possibly misfolded protein for degradation (48). Our results suggest that co-expression of DHHC1 with A457P increases the fragmentation of membrane-bound NET but does not specify which membrane they are bound. One explanation for this may be due to mannose-6-phosphate (M6P) dependent and M6P-indpendent trafficking for degradation (49,50). Further studies on the nature of this fragmentation including the pathway and its final destination may be areas to follow in the pursuit of uncovering NET processing and involvement in the pathogenesis of POTS.

**FIGURE 10.**
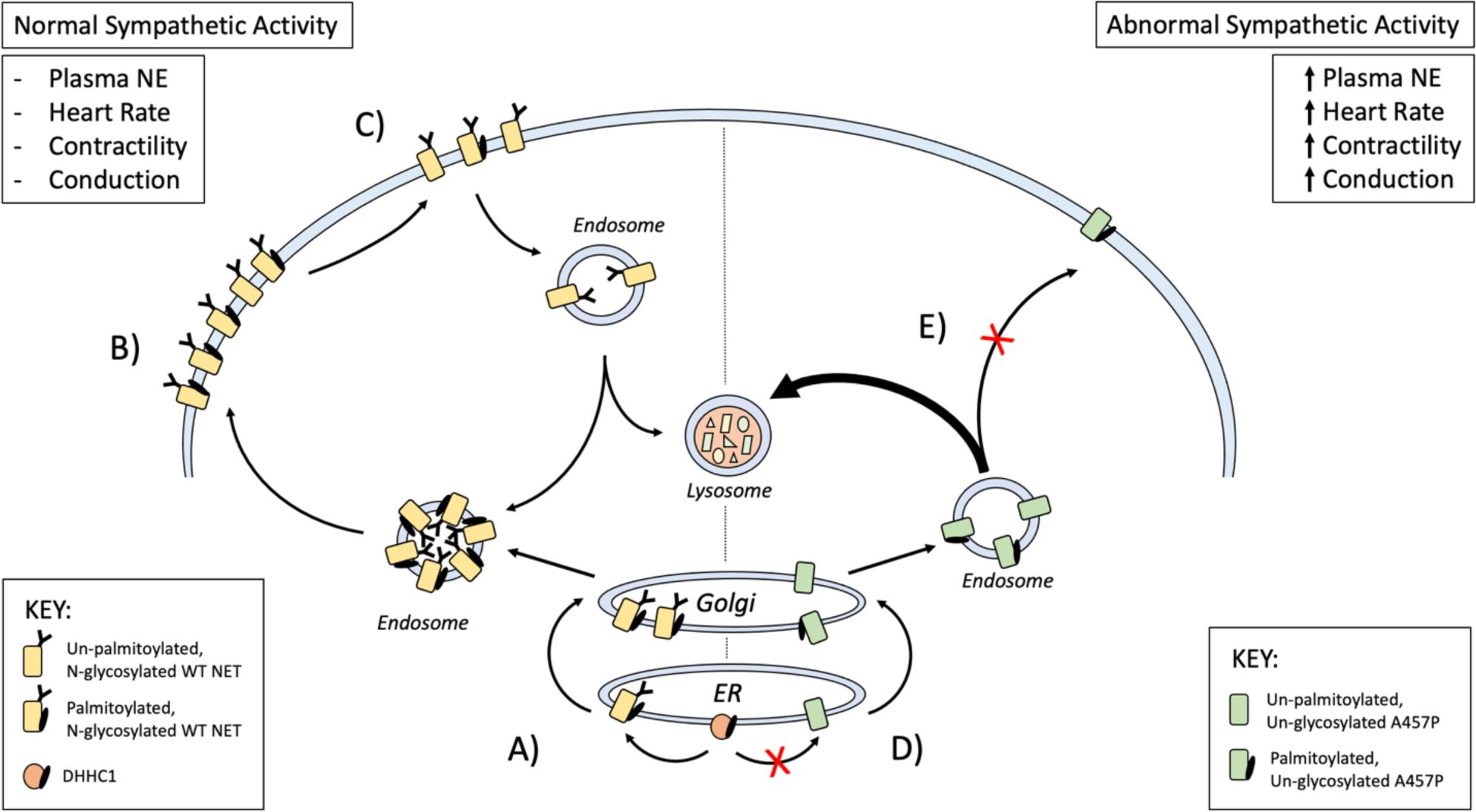
Model for NET palmitoylation and potential dysregulation in POTS. (***A***) Enhancing palmitoylation of mature, N-glycosylated, WT hNET by DHHC1 facilitates ER-Golgi processing and increases surface and total NET levels **(*B*)**. (***C***) Acute inhibition of NET palmitoylation by 2BP dramatically decreases NET surface expression, transport capacity, and total protein levels that continues with long-term inhibition. (***D***) The A457P hNET variant exhibits a large reduction in total mature NET protein levels. Following co-expression of DHHC1, a resultant increase in membrane-bound NET fragments suggests that palmitoylation occurs in sequence with N-glycosylation in an unknown ER processing mechanism, with N-glycosylation potentially being required for palmitoylation-mediated NET stabilization and trafficking or vice versa. (***E***) Inhibition of palmitoylation either through environmental stimuli, pharmacologic strategies, or coding variants leads to NET destabilization and surface trafficking, with a resultant increase in lysosomal degradation. Ultimately, decreased total and surface NET result in poor sympathetic control and pathologic sequela like excessive plasma norepinephrine, tachycardia, and syncope.

Since we are unable to reliably determine the level of palmitoylation in the A457P variant we do not know if DHHC1 co-expression increases its palmitoylation. It is possible that increased DHHC1-mediated palmitoylation of other proteins involved in the trafficking of misfolded integral membrane proteins for lysosomal degradation are responsible for the increase in NET fragments observed with DHHC1 co-expression. Overall, it seems likely that palmitoylation is a single step in a series of post-translational modifications that are required for appropriate NET processing and trafficking. This hypothesis is further strengthened by evidence of other transmembrane proteins undergoing similar post-translational mechanisms. CFTR is processed in a similar manner, with ER-palmitoylation of WT and/or F508del CFTR leading to Golgi sequestration, suggesting that another post-translational event is necessary for complete maturation and surface recruitment of the transporter. This is supported with additional evidence that the large conductance calcium-activated “big” potassium (BK) channel undergoes a similar mechanism, with unpalmitoylated BK channels being withheld from surface recruitment at the trans-Golgi network (51). Demonstrated by our working cellular model in Figure 10, palmitoylation appears to control NET transport capacity through a trafficking dependent mechanism, with changes in transport capacity mirrored by surface density and not changes in intrinsic kinetic parameters. However, it is not known if this process is involved in surface recruitment, internalization, sub-cellular sequestration, or another mechanism. Collectively, it appears that many proteins are regulated in a complex, and sometimes sequential, manner by post-translational modifications that serve to fine-tune its processing, trafficking, and kinetic parameters.

Ultimately, these results outline a novel-mechanism in the regulation of NET trafficking, expression, and activity, and that disruption of this process likely contributes to the pathogenesis of POTS. The role of palmitoylation in regulating NET biogenesis and trafficking has not been previously reported and provides a new collection of evidence in support of palmitoylation-based diseases. Dysregulation of palmitoylation is found in numerous diseases including X-linked intellectual ability (52), Huntington’s (53), Alzheimer’s (54), and amyotrophic lateral sclerosis (55). These new findings suggest that idiopathic POTS may be due to similarly dysregulated processes and may provide opportunities for therapeutic development. It is possible that rescue strategies of NET biogenesis mediated through palmitoylation could serve as a plausible therapeutic option for autonomic disorders like POTS. Nonetheless, additional studies investigating which DHHCs palmitoylate NET, the potential reciprocal relationships with other modifications (34), and synergistic processes are needed to create a more clear picture of NET regulation and pathogenic processes.

## ACKNOWLEDGEMENTS

Funding: NIDA grant 2R15DA031991-02A1, NSF grant 1852459, and P20 GM103442 (IDeA) from INBRE of NIGMS and UND SMHS. We thank Dr. Masaki Fukata, National Institute for Physiological Sciences, Japan, for the generous gift of DHHC coding plasmids.

## AUTHOR CONTRIBUTIONS

C.R.B conceptualized, performed the experiments, and created the figures in this study under the supervision of J.D.F. C.R.B wrote and edited the original version of the manuscript and J.D.F contributed to conceptualization and editing of the manuscript’s final version.

